# Physical activity shapes the intestinal microbiome and immunity of healthy mice but has no protective effects against colitis in MUC2^-/-^ mice

**DOI:** 10.1101/2020.05.24.113290

**Authors:** Mehrbod Estaki, Douglas W. Morck, Candice Quin, Jason Pither, Jacqueline A. Barnett, Sandeep K. Gill, Deanna L. Gibson

## Abstract

The interactions among humans, their environment, and the trillions of microbes residing within the human intestinal tract form a tripartite relationship that is fundamental to the overall health of the host. Disruptions in the delicate balance between the intestinal microbiota and their host immunity are implicated in various chronic diseases including inflammatory bowel disease (IBD). There is no known cure for IBD, therefore, novel therapeutics targeting prevention and symptoms management are of great interest. Recently, physical activity in healthy mice was shown to be protective against chemically-induced colitis, however the benefits of physical activity during or following disease onset is not known. In this study, we examine whether voluntary wheel running is protective against primary disease symptoms in a mucin 2 deficient (*Muc2^-/-^*) life-long model of murine colitis. We show that 6 weeks of wheel running in healthy C57BL/6 mice leads to distinct changes in fecal bacteriome, increased butyrate production, and modulation in colonic gene expression of various cytokines, suggesting an overall primed anti-inflammatory state. However, these physical activity-derived benefits are not present in *Muc2^-/-^* mice harboring a dysfunctional mucosal layer from birth, ultimately showing no improvements in clinical signs. We extrapolate from our findings that while physical activity in healthy individuals may be an important preventative measure against IBD, for those with a compromised intestinal mucosa, a commonality in IBD patients, these benefits are lost.

## Introduction

Inflammatory bowel diseases (IBD) encompassing Crohn’s disease (CD) and Ulcerative colitis (UC) are idiopathic, relapsing chronic diseases characterized by chronic inflammation of the gastrointestinal tract. While pathology varies between UC and CD, both burden patients with common debilitating clinical symptoms such as diarrhea, rectal bleeding, abdominal pain, and weight loss. The etiology of IBD is not known, however a combination of genetic, immunological, and environmental factors is implicated in its development. Most recently, the contribution of the intestinal microbiota in IBD pathogenesis has risen as an active area of research (1). For example, IBD patients have reduced gut microbial diversity (2) and are more likely to have been exposed to antibiotics in 2-5 years preceding their diagnosis (3). In animal models, mice genetically predisposed to colitis (IL-10^-/-^) are resistant to disease onset while kept under germ-free conditions, however clinical signs instigate immediately following exposure to microbes (4).

With incidence of IBD and its burdens rising globally (5), there is an increasing demand for novel therapeutics. Physical activity (PA) has been proposed as both a primary and an adjunct therapy for prevention and treatment of various chronic diseases due to its well documented ability to ameliorate low-grade systemic inflammation (6). Most recently, IBD has been marked as a potential new candidate (7) to yield benefits from regular PA. Studies of PA in rodents have shown attenuated clinical signs of chemically-induced colitis (8–11) that appear to be dependent on the colitis model and type of PA. These studies, however, only assess the role of PA as a preventive measure leading up to induction of acute colitis via a chemical toxin. As so, the potential benefits of PA succeeding or during disease onset is not known. In this study we aimed to address this knowledge gap by utilizing the mucin 2 knock-out (*Muc2^-/-^*) mouse model of chronic colitis.

The human intestinal tract is continuously exposed to the trillions of microbes residing within the mucosal layer of the lumen. Under homeostatic conditions, these microbes are tolerated by the host as they provide essential functions such as digestion of complex carbohydrates, protection against enteric pathogens, and production of beneficial short-chain fatty acids (SCFA), to name a few. Separating the luminal microbes from intestinal epithelial cells (IEC) is a mucus bilayer largely composed of the highly glycosylated protein MUC2. In the colon, the loosely structured outer mucus layer allows for colonization of microbes in a nutrient-rich environment, while the dense inner layer segregates them from the IEC (12). Inflamed intestinal tissues of UC patients commonly display structural defects or thinning of this mucus layer (13), leading to excessive exposure of microbial antigens to the host cells, prompting a chronic state of inflammation and apoptosis leading to further loss of IEC integrity and thus further exposure and injury. *Muc2^-/-^* mice or those with missense mutations impairing the release of MUC2, are born with an underlying predisposition to intestinal inflammation that show rapid progression of colitis (14). *Muc2^-/-^* mice generally display early clinical signs of colitis following weaning (~ 1 month) and microscopic tissue pathology as early as two months of age, indicating a moderate-level colitis, reaching high-severity by 4 months (15).

We hypothesized that introduction of *Muc2^-/-^* mice to voluntary wheel running (VWR) immediately following weaning would reduce the severity and delay the onset of clinical signs of colitis. Having recently shown a significant correlation between aerobic fitness and overall microbial diversity and increased butyrate production (16), we hypothesized further that PA-associated protection would be mediated through changes of the intestinal microbiota and their metabolites.

## Methods

### Experimental procedure

To test our primary hypothesis that habitual physical activity can be protective during IBD as a treatment, rather than a preventative therapy as has been previously shown, we chose to utilize *Muc2^-/-^* mice as a life-long model of murine colitis. In our facility, *Muc2^-/-^* are underweight at birth and display early signs of colitis that are mild to moderate in severity up to ~ 3 months of age, at which point they accelerate rapidly reaching high severity by 4 months. We therefore designed our experiment to conclude prior to the 3 month time-point under the speculation that higher severity of disease symptoms would preclude the animals from voluntarily running on wheels. Following weaning at 5 weeks of age, animals were randomly assigned to individual cages under one of four groups (n=8 per group): Wild-type (WT) C57BL/6 mice with access to a free running wheel (VWR) or a locked wheel (SED), and *Muc2^-/-^* mice with access to free wheel (MVWR) or a locked wheel (MSED). We chose wheel running as a model of PA over forced exercise, as mice voluntarily run higher total distances on free wheels than when forced on a treadmill (17). Forced exercise can also cause significant stress in rodents (18) and in fact has been shown to exacerbate colitis severity in C57BL/6 mice (9). A six-week VWR intervention period was selected based on previous reports showing this to be sufficient in eliciting significant protection against chemically-induced models of colitis (19, 20). The primary responses of interest during this period were weekly clinical signs of colitis and histopathological disease scores at terminus. Additional responses of interest were changes to intestinal microbial composition and function, immunity, and SCFA production in response to PA. The variables associated with these additional responses are described in detail below. By examining these we aimed to elucidate the mechanisms by which PA recruits protective actions.

### Animals

All procedures involving the care and handling of the mice were approved by the UBC Committee on Animal Care, under the guidelines of the Canadian Council on Animal Care. Four-week-old male wild-type (WT) C57BL/6 mice were purchased from Charles River (Vancouver, CA) and kept under specific pathogen-free conditions. *Muc2^-/-^* mice, generated also on a C57BL/6 genetic background, were bred in house with the founding colonies kindly donated by Dr. Bruce Vallance from the Child and Family Research Institute (UBC Vancouver). All animals were housed in a temperature-controlled room (22 + 2°C) on a 12h light/dark cycle with access to acidified water and irradiated food (PicoLab Rodent Diet 20-5053, Quebec, CA) *ad libitum*. Assignment of mice to experimental groups were carried out using a random number generator immediately preceding individual cage allocation.

### Voluntary wheel running, and food and water intake

The running wheels (Columbus Instruments, diameter 10.16 cm, width 5.1 cm) were mounted to the top of the cage lids and were programmed to record the total number of revolutions at 1 hr intervals for the duration of the experiment. Body weights, food consumption, and water intake were measured weekly at approximately the same time during the light cycle. Food weight measurements consisted of subtracting the week’s remaining pellets on the cage lids and bottoms from that week’s starting weight.

### Tissue collection

For fecal sample collection, mice were kept briefly in isolation in sterile and DNAzap-treated containers until defecation. Collected fecal pellets, which were used for microbiome surveying, were immediately snap-frozen in liquid nitrogen then stored in −80 °C until further analyses. Fecal samples were collected on day 1 immediately following assignment to individual cages, and again on the final experiment day immediately preceding tissue collection. Animals were euthanized by cervical dislocation while under deep isoflurane anesthesia. The cecum was isolated, its content removed, and tissue frozen in liquid nitrogen for further analyses of SCFA composition. Colon tissues were collected as follows: starting from distal end, two consecutive ~1.5 cm sections were collected with the most distal section being fixed in 10% neutral buffered formalin for histopathology and the proximal section was stored in RNAlater (Thermo Fisher Scientific) for use in cytokine gene expression assays. All frozen samples were then stored at −80 °C until further use.

### Clinical and Histopathological Scoring

Disease progression and severity in *Muc2^-/-^* animals was assessed based on an in-house clinical signs scoring system, and represented by a variable we henceforth call “disease score”. Briefly, each animal was graded weekly based on the following: observed behavior from a distance, stool/rectal bleeding, stool consistency, weight loss, and hydration, with each variable being assigned a score of 0-4. Humane endpoint was set as a total cumulative score of ≥12, rectal prolapse, or a weight loss of >20% body weight for 2 consecutive days. No animals reached a humane endpoint in this study.

For histopathological scoring, colon cross-sections were fixed in 10% neutral-buffered formalin at 4°C overnight, washed 3 times with phosphate buffered saline (PBS, pH 7.4), transferred to 70% ethanol and sent for dehydration, paraffin-embedding, sectioning, and hematoxylin and eosin (H&E) staining at Wax-it Histology Services (Vancouver, Canada). Tissue slides were coded throughout the microscopy analyses and investigators scoring histopathology were blinded to the groupings. H&E stained sections were viewed under 200x magnification on an Olympus IX81 microscope and the full image stitched together using MetaMorph^®^ software. Stitched images were imported into ImageJ-version 1.51r (21) for scoring. Disease severity in colonic cross sections from the *Muc2*^-/-^ animals were assessed using a previously described scoring system (22). In brief, a total score was calculated for each mouse using the following criteria

1. *Edema*, as compared to a healthy WT control: 0=no change; 1 =mild (<10%); 2 = moderate (10-40%); 3=profound (>40%)
2. *Epithelial hyperplasia*, average height of crypts as a percentage above the height of a healthy control where 0=no change; 1 = 1–50%; 2=51–100%; 3=>100%
3. *Epithelial integrity*, shedding and shape of the epithelial layer as compared to healthy control where: 0=no change; 1 = <10 epithelial cells shedding per lesion; 2 = 11–20 epithelial cells shedding per lesion; 3=epithelial ulceration; 4=epithelial ulceration with severe crypt destruction
4. *Cell infiltration*, presence of immune cells in submucosa: 0=none; 1 = mild (2-43); 2 = moderate (44-86); 3= severe (87-217).

The resulting variable, henceforth called “histopathological score”, has a maximum value of 13.

### Reverse Transcriptase-qPCR

To identify the potential immunological pathways involved in PA-derived protection, we examined the gene expression of several key immune markers commonly associated with colitis. The mRNA gene expression for tumor-necrosis factor alpha (TNFα), interferon-gamma (IFNγ), resistin-like molecule beta (Relm-β), regenerating islet-derived protein 3 (RegIII-γ), transforming growth factor beta (TGF-β), chemokine C-X-C motif ligand 9 (Cxcl9), and claudin 10 (Cldn10) were measured in colon tissues. Total RNA was purified from tissues using Qiagen RNEasy kits (Qiagen) according to the manufacturer’s instructions with an additional initial bead beating step (3×30 seconds, 30 Hz) on a Retsch MixerMill MM 400 homogenizer. Next, cDNA was synthesized using the iScript cDNA Synthesis Kit (Bio-Rad) in 10 μl reactions. The RNA and cDNA products’ purity and quantity were assessed by a NanoDrop spectrophotometer (Thermo Scientific). The cDNA products were normalized to ~ 40 ng/μl with DNAse free sterile water prior to qPCR reactions.

A total of 10 μl RT-qPCR reactions consisted of: 0.2 μl of each forward and reverse primers (10mM), 5 μl of Sso Fast Eva Green Supermix (Bio-Rad), 3.6 μl DNAse free water, and 1 μl of cDNA template. Reactions were run in triplicates using the Bio-Rad CFX96 Touch thermocycler and analyzed using Bio-Rad CFX Maestro software 1.1 (v4.1). The median quantitation cycle (Cq) value from each sample was used to calculate the 2^−ΔΔCt^ based on the reference gene TATA box binding protein (Tbp). A list of all the primer sets, their melting temperature, efficiencies, and detailed thermocycler protocol used in this study are described in Supplementary Material 6.

### Short-chain fatty acids

SCFAs, a byproduct of microbial fermentation, are an essential component of a healthy gut environment (reviewed in (23)). They not only serve as a primary food source for the colonocytes, but have immunogenic properties that, in concert with the host immunity, are integral in maintaining gut homeostasis. We previously showed that in healthy humans, cardiorespiratory fitness was positively correlated with fecal butyrate (16), a SCFA with known anti-inflammatory properties in the gut (24). We therefore hypothesized that SCFA profiles of VWR mice would differ from SED, favoring the production of beneficial butyrate that may be involved in protection against colitis. We therefore analyzed SCFA (acetic, propionic, heptanoic, valeric, caproic, and butyric acids) in cecal tissues by gas chromatography (GC) as described previously in (25). In brief, ~50 mg of stool was homogenized with isopropyl alcohol, containing 2-ethylbutyric acid at 0.01 % v/v as internal standard, at 30 Hz for 13 min using metal beads. Homogenates were centrifuged twice, and the cleared supernatant was injected to Trace 1300 Gas Chromatograph, equipped with Flame-ionization detector, with AI1310 autosampler (Thermo Fisher Scientific) in splitless mode. Data was processed using Chromeleon 7 software. Half of the cecal tissue was freeze dried to measure the dry weight, and measurements are expressed as μmol/g dry weight (d.w).

### DNA extraction and 16S rRNA amplicon preparation

The effects of PA on the intestinal microbiome has recently risen as an area of great interest (Reviewed in (26)). To date, several studies in mice have shown significant changes to the microbiome associated with either VWR or forced treadmill running (19, 27–29). Allen *et. al* (30) further showed that transplanting the microbiome of exercised mice into germ-free mice conferred protection against dextran-sodium-sulfate (DSS)-induced colitis, highlighting the importance of PA-derived changes in the microbiome. To examine such potential changes in gut microbiome, we surveyed the fecal microbiome of our mice using high-throughput sequencing. DNA was extracted from fecal samples using the QIAmp DNA Stool Mini Kit (Qiagen) according to the manufacturer’s instructions following 3 x 30 s of beat beating as before. Amplicon libraries were prepared according to the Illumina 16S Metagenomic Sequencing Library Preparation manual. In brief, the V3-V4 hypervariable region of the 16S bacterial rRNA gene was amplified using recommended 341F and 805R degenerate primer sets, which create an amplicon of ~460 bp. Amplicons were purified using AMPure XP beads and adapters and dual-index barcodes (Nextera XT) were attached to the amplicons to facilitate multiplex sequencing. Following a secondary clean-up step, libraries were quality controlled on an Experion automated electrophoresis system (Bio-Rad), and sent to The Applied Genomic Core (TAGC) facility at the University of Alberta (Edmonton, Canada) where they were normalized using fluorometric method (Qubit, Thermo Fisher Scientific) and sequenced using the Illumina MiSeq platform with a V3 reagent kits allowing for 2 x 300 bp cycles.

### Bioinformatics

All bioinformatics processes were performed using a combination of R statistical software (31) and the QIIME 2 platform (32) using the various build-in plugins described below. Demultiplexed sequences were obtained from the sequencing facility and primers removed reads using *cutadapt* (33). Sequences then underwent quality-filtering, dereplication, denoising, merging, and chimera removal using DADA2 (34). The output of this process is a feature table of amplicon sequence variants (ASV) that is a higher resolution analogue of traditional OTU tables. To aid in removal of non-specific host contaminants, a positive filter was applied to all reads using the latest available Greengenes (13_8) (35) database (clustered at 88% identity). All ASVs were searched against the reference reads using VSEARCH (36) and any that did not match the reference sequences at a minimum of 70% identity similarity at 70% alignment were discarded. For analyses encompassing phylogenetic information, a phylogenetic tree was constructed using a SATé-enabled phylogenetic placement (SEPP) technique as implemented in the *q2-fragment-insertion plugin* (37) using a backbone tree build based on the SILVA (128) database (38). Taxonomic classification of the ASVs were carried using IDTAXA (39). It has been proposed that the functional repertoire of the gut microbiota is more sensitive to perturbation than taxonomic changes, and therefore may be crucial in identifying underlying physiological signals (40). To predict the functional potential and phenotype of the microbiome, we used BugBase (41) which utilizes PICRUSt’s (42) extended ancestral-state reconstruction algorithm for metagenome composition prediction. As these tools require sequences to be classified against the Greengenes taxonomy assignments, we used VSEARCH to pick closed-reference OTUs from our denoised feature table at 97% similarity threshold against the 99% identity clustered Greengenes database.

### Statistical Analyses

All statistical analyses were performed using R version 3.5.1 unless stated otherwise. During the 3rd week of the experiment, the VWR animals were unintentionally exposed to 3 days of irregular light-dark cycles as a result of an electrical malfunction with the lighting in the animal room. While the exact nature of this disruption is not known, the wheel running data during this period suggests a period of reduced activity. The issue was resolved by the 3rd day and the animals did not display any signs of stress or irregular behavior; we therefore consider this to be of minimal impact to the experiment. However, as a precaution, we chose to analyze the data as a 4 x 1 (*groups*) factorial design rather than 2 x 2 (*activity x genotype*) as we could not definitively eliminate the possibility that wheel running in this group was impacted by the brief interruption.

### Wheel running

To determine whether WT and *Muc2^-/-^* ran similar distances throughout the experiment, we first analyzed total weekly distances (km) run by each group across the 6 weeks time using linear mixed-effects regression (LMER) using the *lme4* package with individual animals set as the random effect and *groups* as the fixed effects. Homoscedasticity and linearity of the models were assessed using diagnostic plots of the residuals.

### Body weights and food/water intake

To monitor overall behavioral changes of mice as a result of PA between WT and *Muc2^-/-^* mice, we examined weekly body weights, food and water intake across the 6 weeks. To account for natural differences in starting body weights, total weight gained relative to starting body weights was calculated each week. Body weight, food and water intake across the 6 weeks were each assessed separately using a repeated measures LMER with *time* coded as a random effect and *groups* as a fixed effect. A Tukey HSD post-hoc test with the Benjamini-Hochberg (BH) P adjustment method was used when an overall significance (set as P<0.05) in the models were detected.

### Clinical and Histopathological Scoring

We used a cumulative link model (CLM) with a logit link to evaluate whether the disease score, and separately, the histopathological score differed among treatment groups. This proportional odds type test is more appropriate for ordinal data than classic linear regressions. For clinical scores, the model included *time* and *groups* as the fixed effects, and individual animal ID as the random effect. For histopathological score, the total average score of the MSED and MVWR groups were separately analyzed using the same method, but without the *time* random effect. We implemented the analyses using the *ordinal* R package.

### Colon mRNA gene expression

To test whether the expression of colonic mRNA genes differed across groups, we first explored the overall abundance of all surveyed genes simultaneously using an ordination method. The Euclidean distances of Hellinger-transformed relative gene expression values were ordinated onto a principal component analysis (PCA) plot. When a clear clustering was observed based on group assignments, differences in variance across these groups were assessed using a permutational multivariate analysis of variance PERMANOVA test using the *vegan* package, and pairwise differences calculated using *pairwiseAdonis* with BH adjustment for multiple testing. For differential abundance testing of each cytokine, a multivariable generalized linear model (GLM) test was carried out using the *mvabund* package (43). This fits separate GLMs to each cytokine while accounting for non-independence and adjusting for multiple testing. The negative binomial distribution assumption was selected for the model and the mean-variance plot was used to assess the model fit. A Kruskal-Wallis post-hoc test was carried on individual genes when significance was detected in the overall model. Pairwise comparisons across groups were carried out using Conover’s test for multiple comparisons within the *PMCMRplus* package.

### Short-chain fatty acids

Similar to the cytokine data analysis, to test the differences in abundance of SCFAs across groups, concentrations of various cecal SCFA were assessed using a multi-GLM test. Post-hoc tests were carried out on individual SCFAs identified as significant from the univariate results from the global model.

### Microbial Analysis

Following our previous observation in humans that showed distinct microbial community characteristics and metagenomic functions associated with higher cardiorespiratory fitness, we evaluated whether similar patterns emerged in mice. Community structural patterns of fecal bacteria across samples (β diversity) were explored using the *q2-DEICODE* plugin (44). DEICODE is a compositionally aware method that utilizes a form of robust Aitchison distances to create a species abundance distance matrix of ASVs which can then be projected onto a PCA biplot. We visualized this using the Emperor interactive graphic tool (45). To reveal possible group differences, a PERMANOVA (46) test was conducted on all groups across time. Pairwise testing was then followed using a Kruskal-Wallis test with a BH adjustment to control for false discovery rate (FDR).

The overall within-sample diversity (α diversity) for each sample was estimated based on the species richness, Simpsons, and Shannon indices using the *DivNet* package (47). For each group, the difference between a sample’s week 6 and week 0 diversity score was calculated and used to determine whether those changes differed from zero (Wilcoxon test) as well as other groups (ANOVA).

Differential abundance testing of individual taxa was performed using the *CornCob* package (48).

BugBase was used to determine high-level phenotypes of bacterial communities based on the following default traits: Gram negative vs. Gram positive, biofilm forming, mobile element containing, oxidative stress tolerance, pathogenic potential, and oxygen utilizing. Pre- and post-treatment differences in relative abundances of these elements were tested in each group using a Kruskal-Wallis test with Benjamini-Hochberg adjustment of P values to control FDR.

## Results

### Wheel running

For unknown reasons, one animal from each group did not run on the wheels and so were excluded from further analyses. The WT group ran an average (SD) of 46.6 (18.4) km in total throughout the 6 weeks, while the *Muc2^-/-^* animals ran slightly less at 40.7 km (21.5) which correspond to ~ 1.3 and 1.1 km/day, respectively. While the WT showed a general trend towards more wheel running, the differences were not statistically significant (Figure 1A) likely due to the highly variable nature of running data.

**Figure 1.**
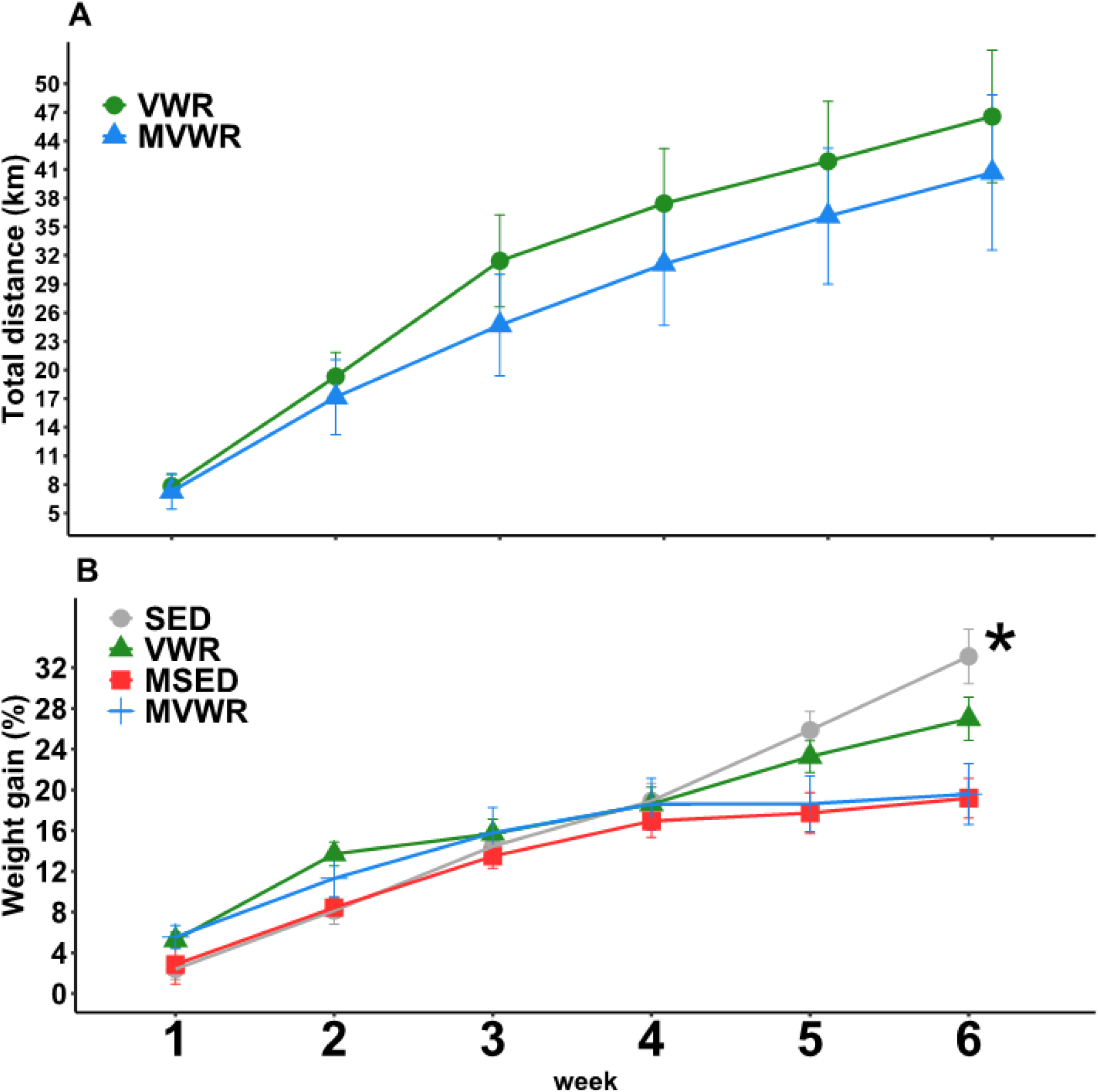
Weekly measures of relative weight gain and wheel running. Longitudinal measurements of A) average accumulated distance ran, B) relative weight gain compared to week 0. Linear mixed models were used with *week* and *animals* set as random effects. There were no significant effects of wheel running in either wheel running or weight gain. * indicates a significant (P<0.05) main effect between genotypes.

### Body Weights, food and water intake

Weight gain was not significantly different across activity levels, however as expected *Muc2*^-/-^ mice gained less weight throughout the 6 weeks (Figure 1B). The mean (± SE) total weight gain of each group was: SED 33.12 ± 2.66 %, VWR 26.98 ± 2.14%, MSED 19.18 ± 1.96%, and MVWR 19.59 ± 2.99% relative to their starting body weights in grams. By the final week, VWR animals had gained ~ 6% less total weight compared to their SED counterpart (P=0.09). Food intake was not statistically different between groups across the 6 weeks (Supplementary Material 1A). *Muc2^-/-^* mice drank significantly more water than WT animals (Beta coefficient (B): 5.4, P<0.001) throughout the 6 weeks. Wheel running was associated with increased water intake in WT (B: 5.3; P<0.01) and to a lesser extent in *Muc2^-/-^* mice (B:1.9; P<0.86) (Supplementary Material 1B).

**Supplementary material 1.**
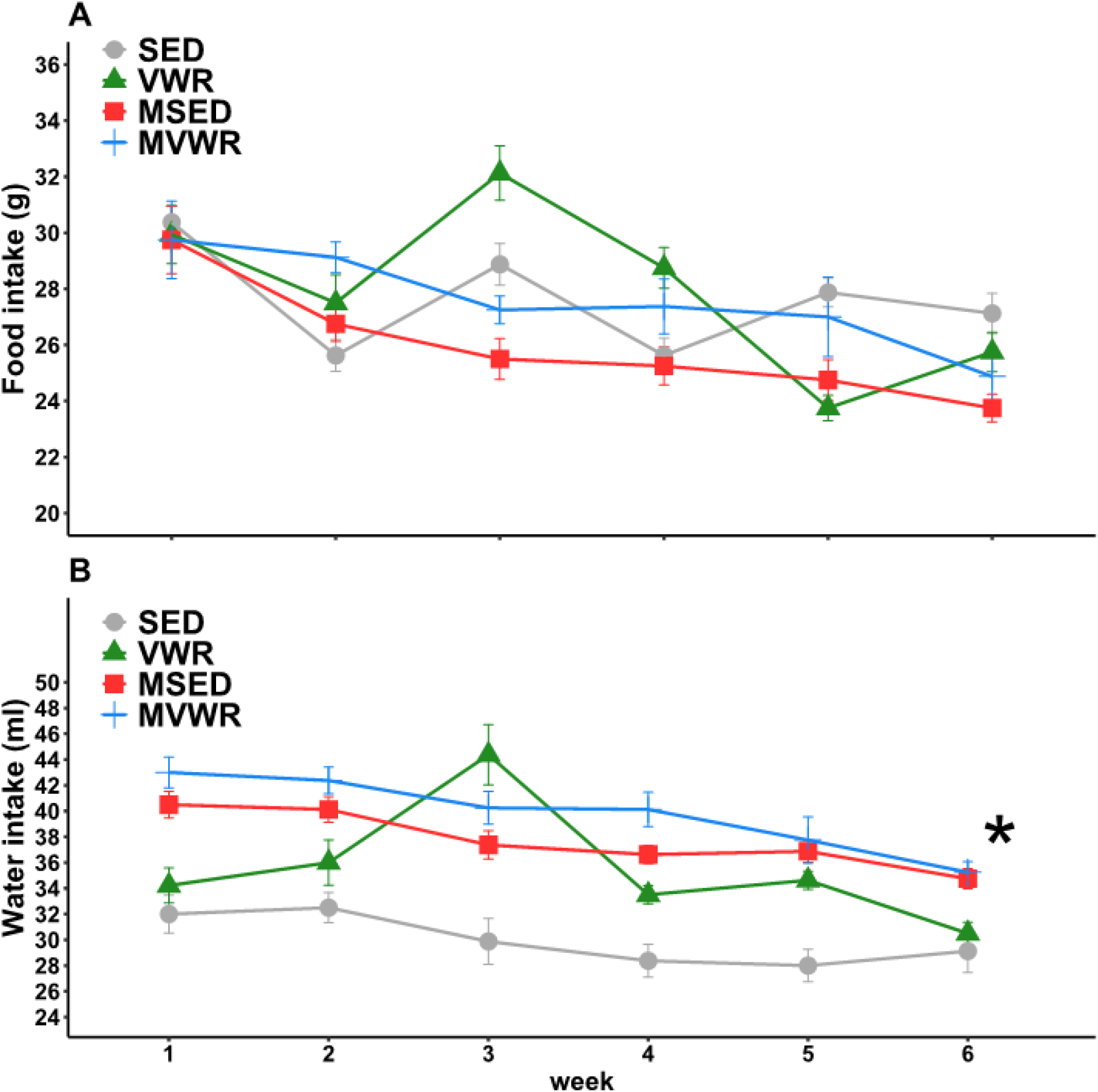
Weekly measures of food and water intake. Longitudinal measurements of A) average weekly food intake, and B) average weekly water intake. Linear mixed models were used with *week* and *animals* set as random effects. There were no significant effects of wheel running in either food or water intake. * indicates a significant (P<0.05) main effect between genotypes.

### Histopathological and clinical scores

We found a modest but significant difference in disease score between MVWR and MSED groups (Coefficient estimate (CE): −1.67; P<0.01) across the 6 weeks, indicating reduced clinical signs in the running animals. However, post-hoc tests carried out each week showed no significant difference between groups (Figure 2A). The differences between groups appear to increase with time, with the largest difference appearing at week 6 (CE: −2.0, P=0.063). Histopathological scores based on H&E sections showed no differences among the groups (Figure 2B).

**Figure 2.**
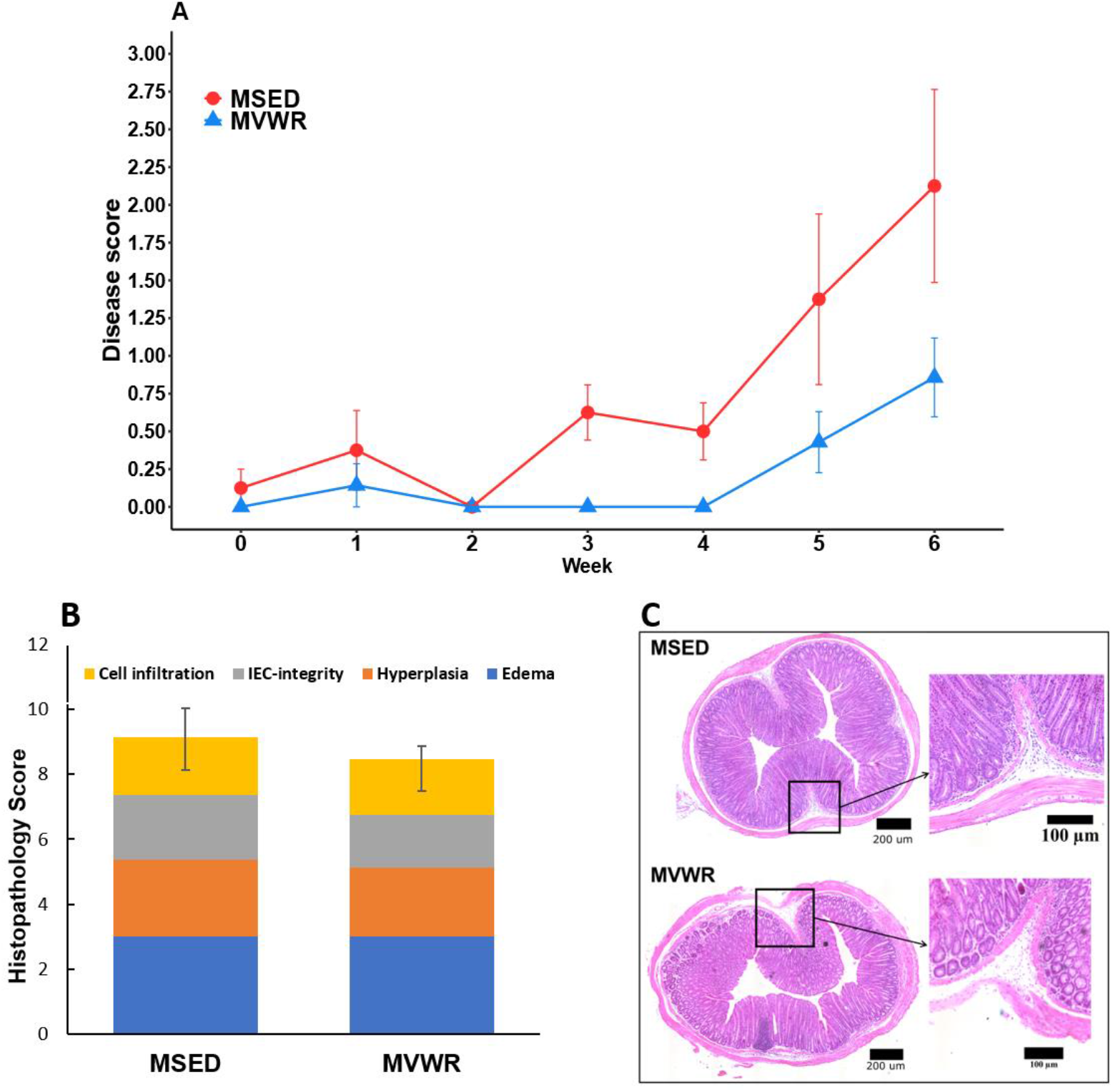
Assessment of severity of colitis signs in *Muc2^-/-^* mice. Comparison of A) clinical disease scores across 6 weeks, and B) histopathological scores in Muc2^-/-^ animals at terminus. C) Representative colon images of H&E stained sections from MSED and MVWR mice. No significant differences were observed between groups in either measurements. Values are shown as means ±SE.

### Short-chain fatty acids

The results of the GLM indicated a significant group effect (Dev: 14.83, P<0.01) and the univariate tests showed significant differences in acetate, propionate, butyrate, and valerate across groups. The results of the post-hoc analyses on these SCFAs and total SCFA are shown in Figure 3. Total SCFA concentration was significantly higher in VWR mice than all other groups, while SED mice had similar total SCFA to both *Muc2^-/-^* groups. VWR mice also had significantly higher total acetate and butyrate than all the other groups and higher propionate than SED. Overall, the major difference between *Muc2*^-/-^ and WT animals was the significantly reduced levels of butyrate in *Muc2*^-/-^ mice and inversely, higher levels of propionate. Valerate, caproate, and heptanoate were similar across all groups. In terms of relative abundances, the main differences between *Muc2*^-/-^ and WT were the higher propionate and lower butyrate proportions in *Muc2^-/-^* animals. Importantly, the proportion of butyrate in VWR mice (~12 %) was significantly higher than those in SED (~7.9 %).

**Figure 3.**
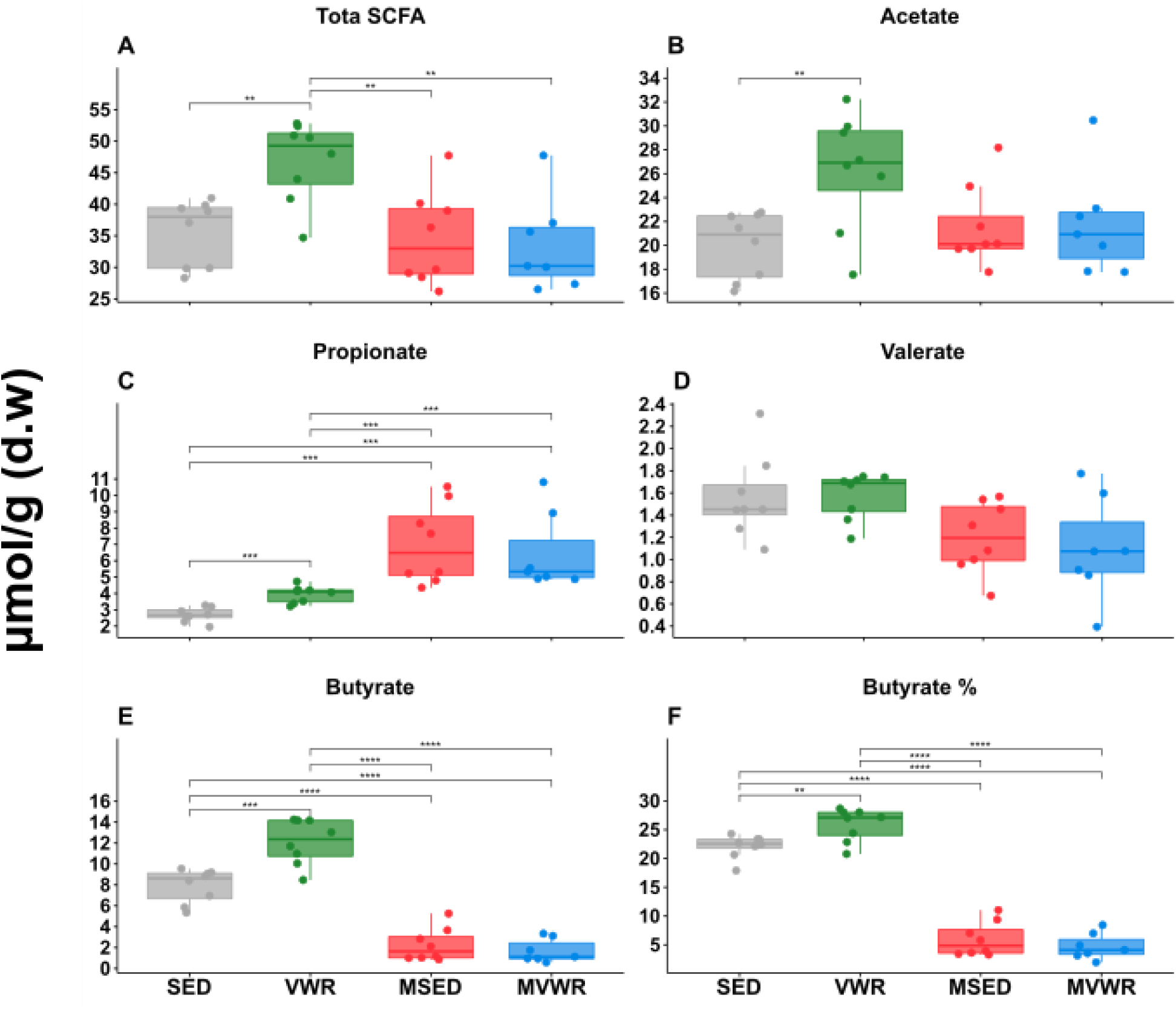
Cecal SCFAs composition. Cecal tissues analyzed for SCFAs composition using gas chromatography. The bottom and top of boxes are the first and third quartiles, the middle band inside the boxes is the median, the whiskers contain the upper and lower 1.5 interquartile range (IQR). * denotes significantly different pairs (adjusted P<0.05)

### Colon mRNA gene expression

The exploratory PCA plot showed clear separation of the WT vs. *Muc2^-/-^* animals along the first principal component (PC1) axis which accounted for 41.7% of the variation (Supplementary Material 2). Further clustering between the SED and VWR groups but not between MSED and MVWR was observed along PC2, which accounted for an additional 17.7% of the variance. The result of the PERMANOVA test confirmed these observations revealing a clear separation amongst groups (F, 10.513; P <0.01). The pairwise comparison test shows a statistically significant separation between all pairs except between MSED and MVWR. The global multi-GLM model showed a significant difference (adjusted P = 0.001) across groups with the univariate tests showing a significant difference in all genes across groups. Notably, VWR mice had significantly lower TNF-α, TGF-β, IFN-γ, and RegIII-γ compared to SED mice (Figure 4); no changes were detected between MVWR and MSED animals. *Muc2^-/-^* animals had increased concentrations of IL-10, RELM-β, CXCL9, RegIII-γ, and TNF-α compared to their WT counterparts.

**Figure 4.**
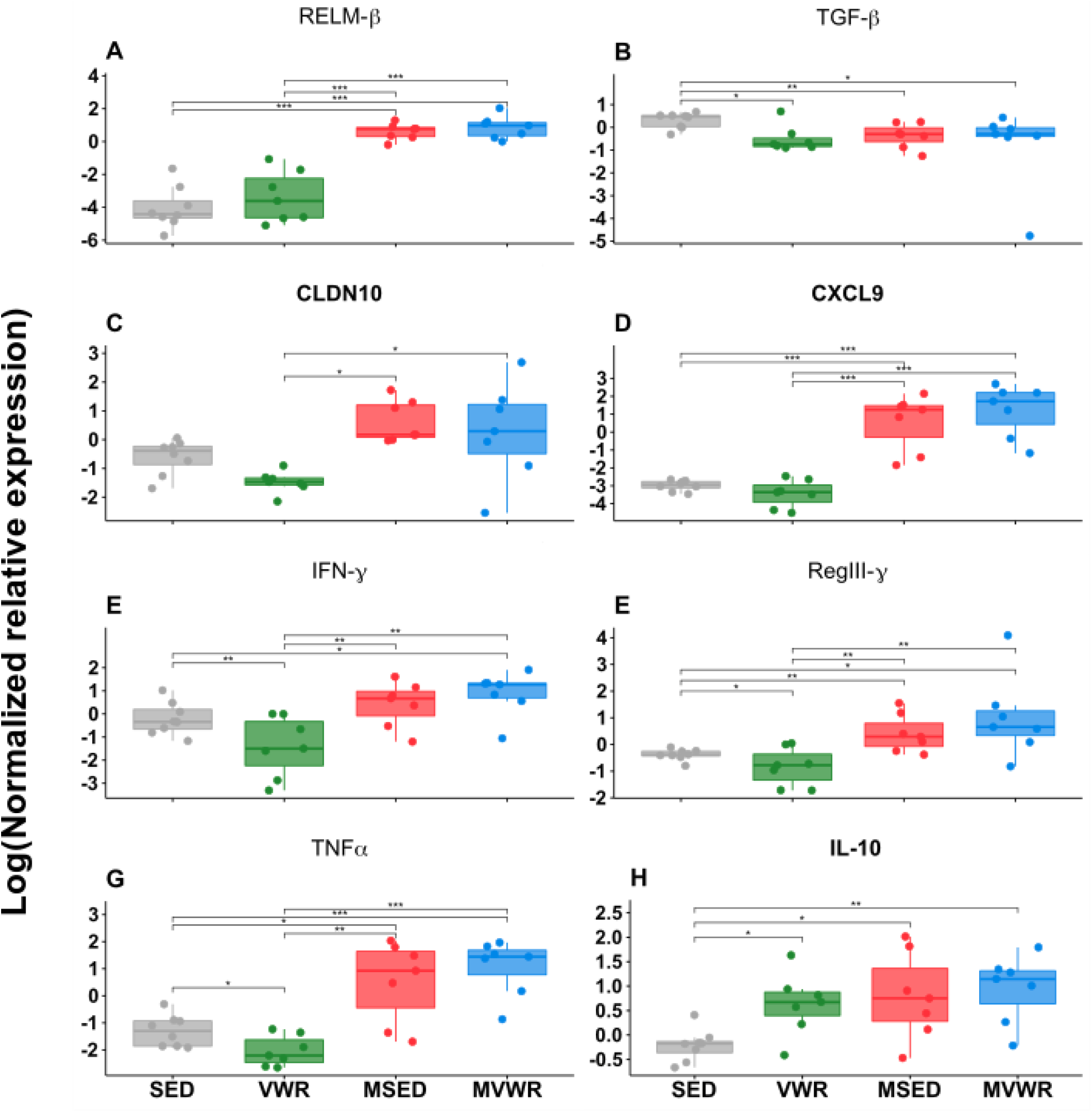
Colonic mRNA gene expression. The relative mRNA gene expression of selected pro- and-anti-inflammatory mediators in colon. The bottom and top of boxes are the first and third quartiles, the middle band inside the boxes is the median, the whiskers contain the upper and lower 1.5 interquartile range (IQR). * denotes significantly different (adjusted P<0.05)

**Supplementary Material 2.**
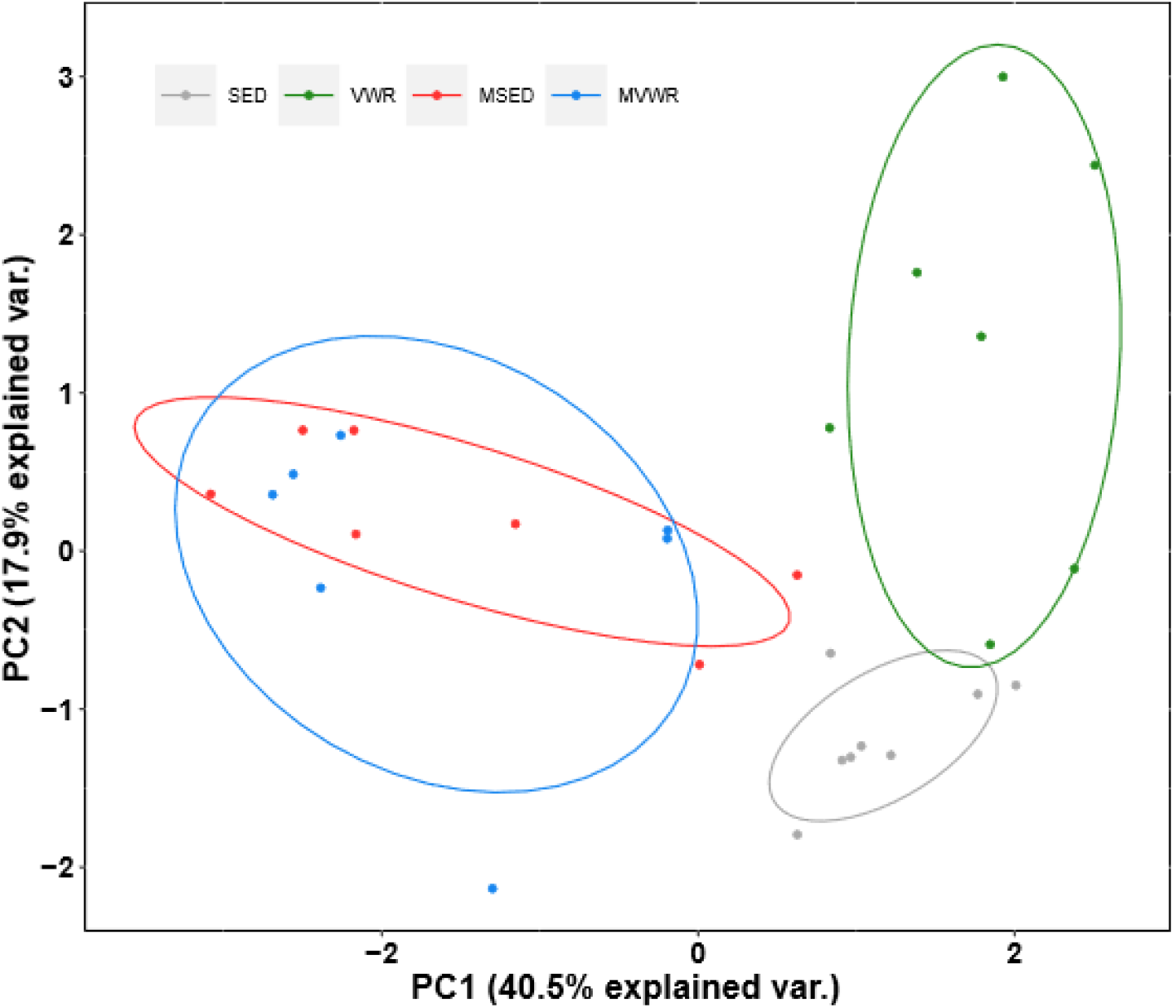
PCA of colonic mRNA gene expression. The Hellinger-transformed Euclidean distances of cytokine abundance data are projected onto PCA. Results of the PERMANOVA test showed a significant (F, 10.513; P <0.01), permutation=999) separation between all groups except between MSED and MVWR.

### Bacterial community analysis

Beta diversity: The PERMANOVA test showed a significant difference between *Muc2^-/-^* and WT animals corresponding to clear clustering observed between these groups on the PCA plots (Figure 5A). Importantly however, there was a significant distance between SED and VWR animal clusters prior to treatment assignment. This fact strongly suggests the presence of a batch effect in our experiment which is likely explained by the fact that the VWR animals were purchased at different times compared to the other groups and their microbiome sequenced separately. As batch-effects are a well-known issue in short-read sequencing experiments (49), differences across groups are then likely confounded by this. Therefore, to mitigate this effect, in all subsequent analyses, changes in microbiome are either only compared within the same group across time, or the change within each group is compared to changes in other groups. Pairwise analysis of each group comparing their week 0 to week 6 profiles showed changes in overall microbiome variation in all animals across time. In *Muc2^-/-^* animals, these changes were non-uniform and did not follow a predictable pattern (Figure 5C). WT animals on the other hand showed a clear and unidirectional change (clear clustering) in their overall microbiome (Figure 5B). A significant shift in community structure of both VWR and SED animals based on the Aitchison distances was observed by week 6 (Pseudo F= 75.2; P=0.001, permutations=999), and these changes were significantly different from each other. These shifts suggest a distinct change in the structure of the microbiome as a function of time as well as physical activity in WT animals. We next compared the magnitude of change (delta=βwk6-βwk0) across all groups and found no significant differences (Supplement Material 3) between them.

**Figure 5.**
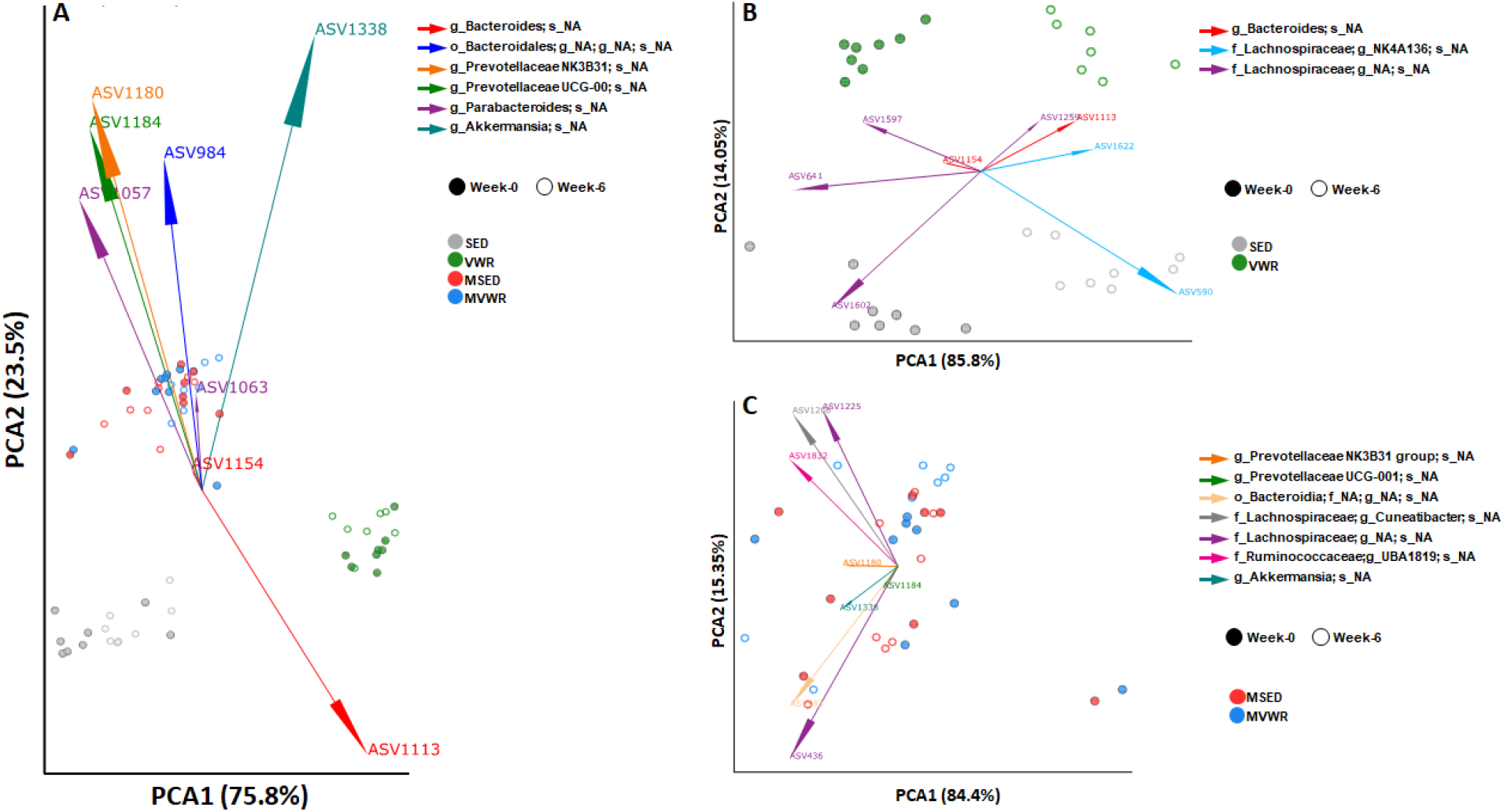
Changes in fecal microbiome composition across time and physical activity. PCA biplot of Robust Aitchison distances visualized where data points represent individual mice colored by their group designation, with spheres corresponding to samples at week 0 and rings representing week 6. The vectors represent the topmost significant ASV loadings driving differences in the ordination space. A) While the *Muc2^-/-^* mice appear to cluster together regardless of activity status, the WT mice appear to be separated at week 0, suggesting the presence of a batch effect, therefore, further analyses are focused on the change across time in each group separately. B) Significant differences were detected as a function of both time and activity in WT mice as determined by PERMANOVA test (pseudo F= 75.2; P=0.001, permutations=999). Pairwise test showed differences between all groups (P<0.001). C) No group differences were detected in *Muc2^-/-^* as determined by a PERMANOVA test (pseudo F= 0.47; P=0.86, permutations=999)

**Supplementary Material 3.**
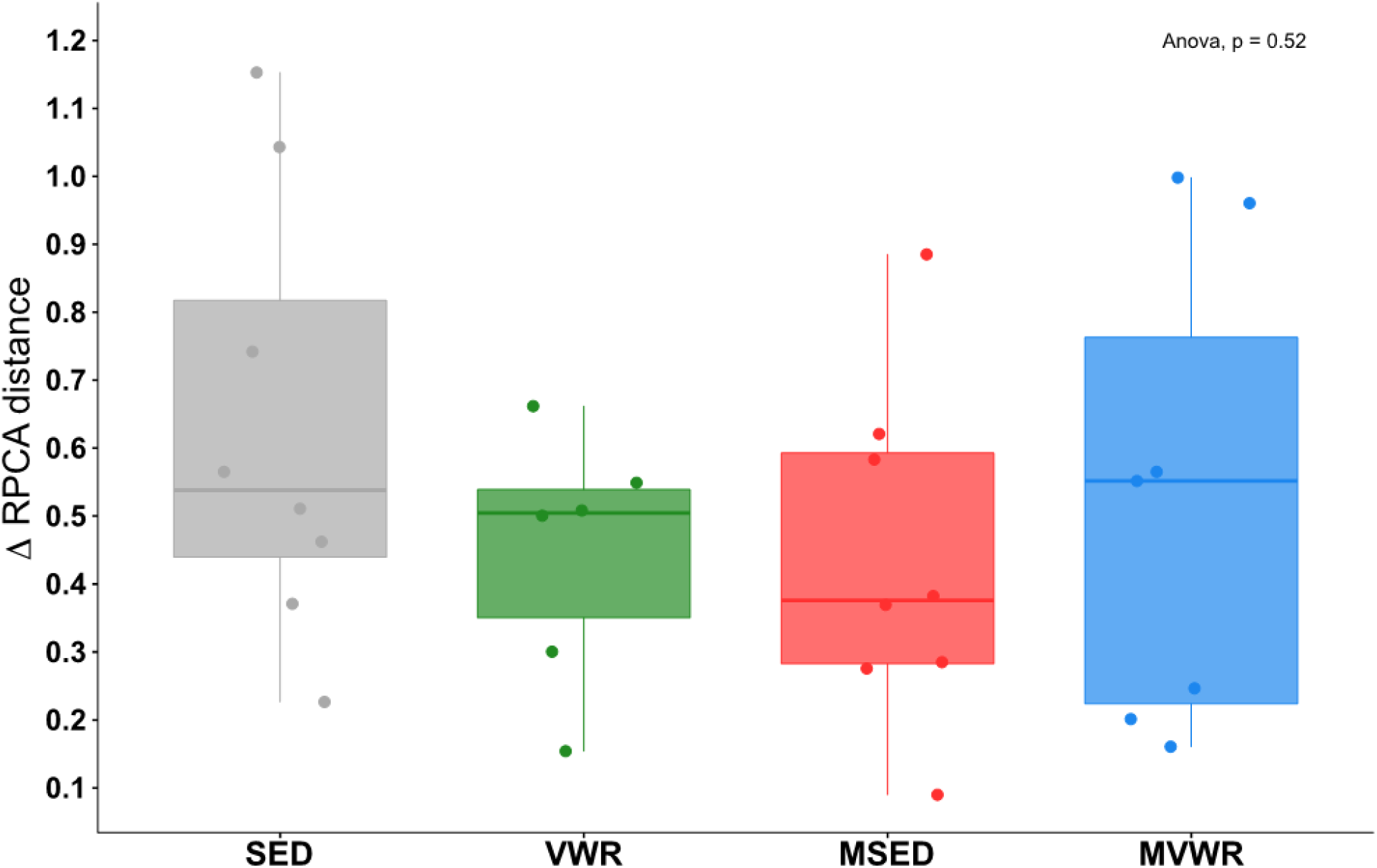
The magnitude of change in beta diversity across time. Robust Aitchison distances were calculated using DEICODE and the change in beta diversity across time within each group was compared to others. All groups demonstrated some change in overall beta diversity across time (values > 0), however the magnitude of change was not different across groups.

Alpha diversity: The difference in change of alpha diversity estimates between week 6 and week 0 showed no significant change across any group (Figure 6A-C). The WT mice had a significantly higher (P < 0.001) overall diversity than *Muc2^-/-^* animals in all examined diversity indexes: species richness, Shannon (Figure 6D), and Simpson index.

**Figure 6:**
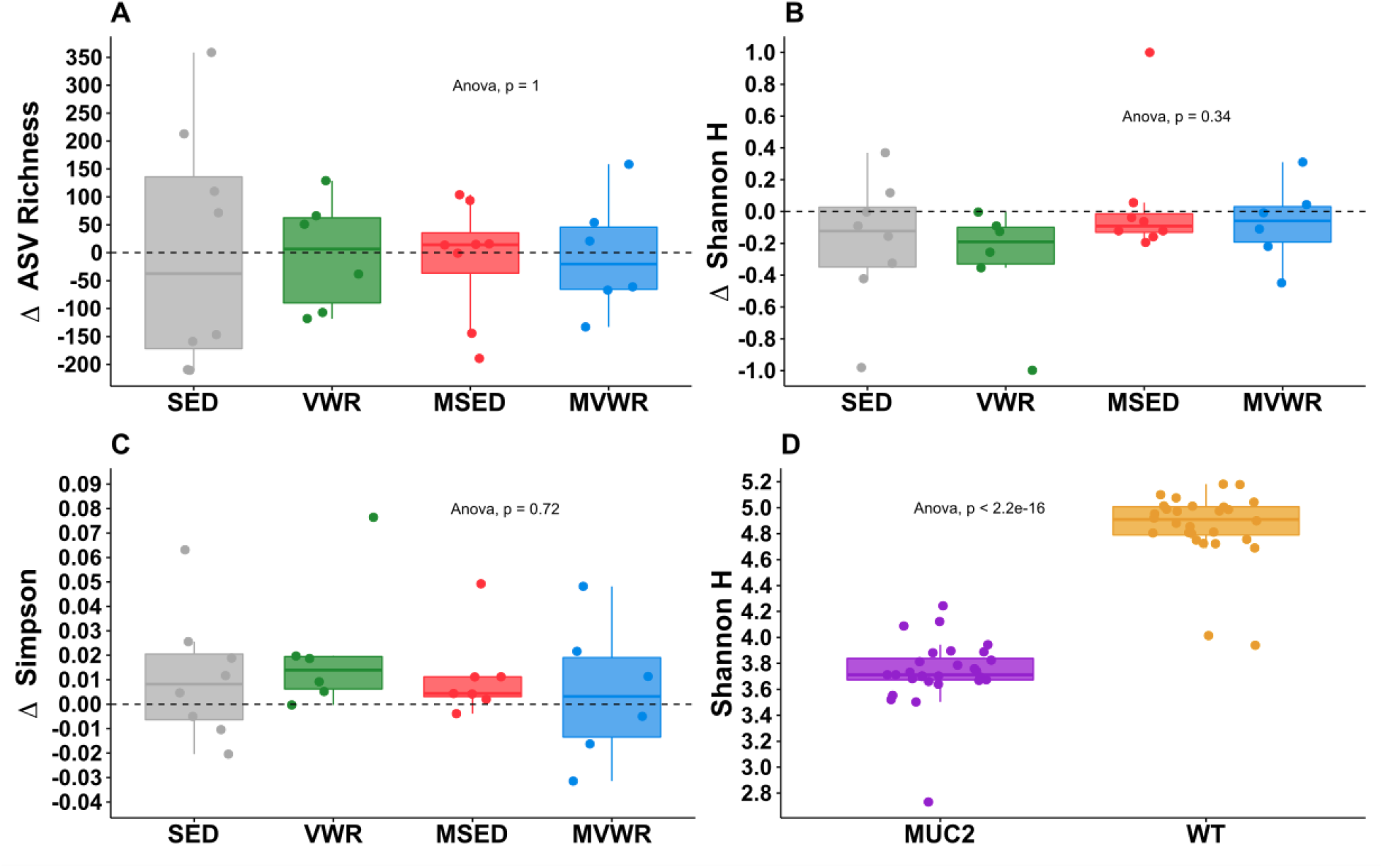
Change in alpha diversity following 6 weeks of voluntary wheel running. Values are differences between week 6 and week 0. Panels A-C, each group’s change in alpha diversity after 6 weeks was compared to its own week 0 values, as well as to other groups. No differences were observed across any groups. D) The overall Shannon diversity of WT mice is significantly higher than those of *Muc2^-/-^* mice

Differential abundance testing: Only WT animals showed statistically significant changes in relative abundance of individual taxa across time. In SED animals, 21 significant taxa were changed by week 6, and 20 were different in VWR animals (Figure 7). Of these 41 total changed taxa, only 4 were common across both groups (Supplementary Material 4). Interestingly, the relative abundance of these 4 taxa changed in the same direction, suggesting the possibility of an age-driven effect. These changes were an increase in the genus Ruminiclostridium5, decrease in genera Negativibacillus and Harryflintia, and decrease in the Lachnospiraceae family.

**Figure 7.**
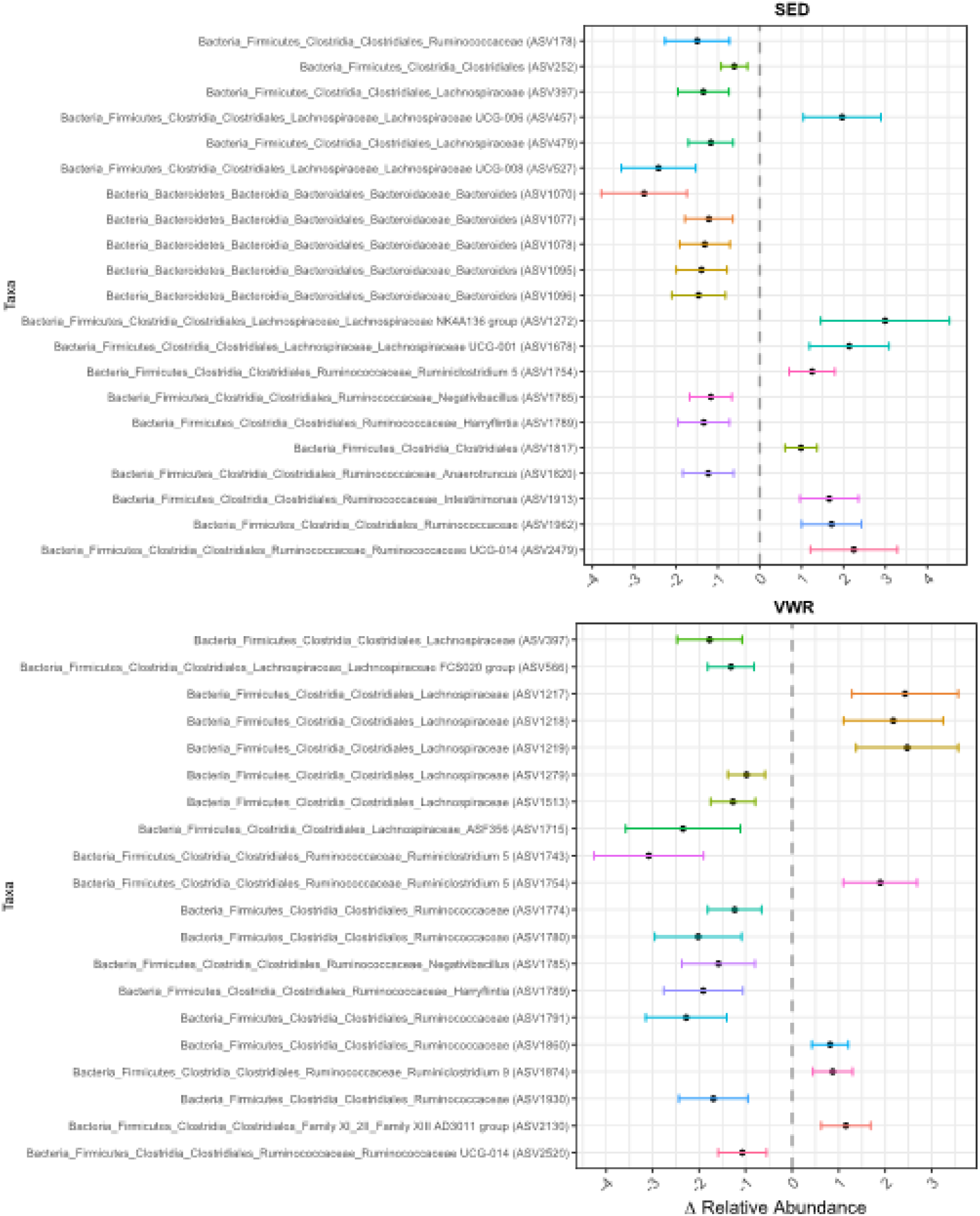
Significant differences in fecal ASVs across time in WT mice. WT mice, but not *Muc2^-/-^*, showed a significant change in relative abundance of several ASVs (threshold set at FDR P <0.01) across time. Different ASVs are changed in VWR compared to SED mice. The points on the plot represent the mean change in relative abundances of each ASVs at week 6 compared to week 0, bars represent the 95% prediction intervals.

**Supplementary Material 4:**
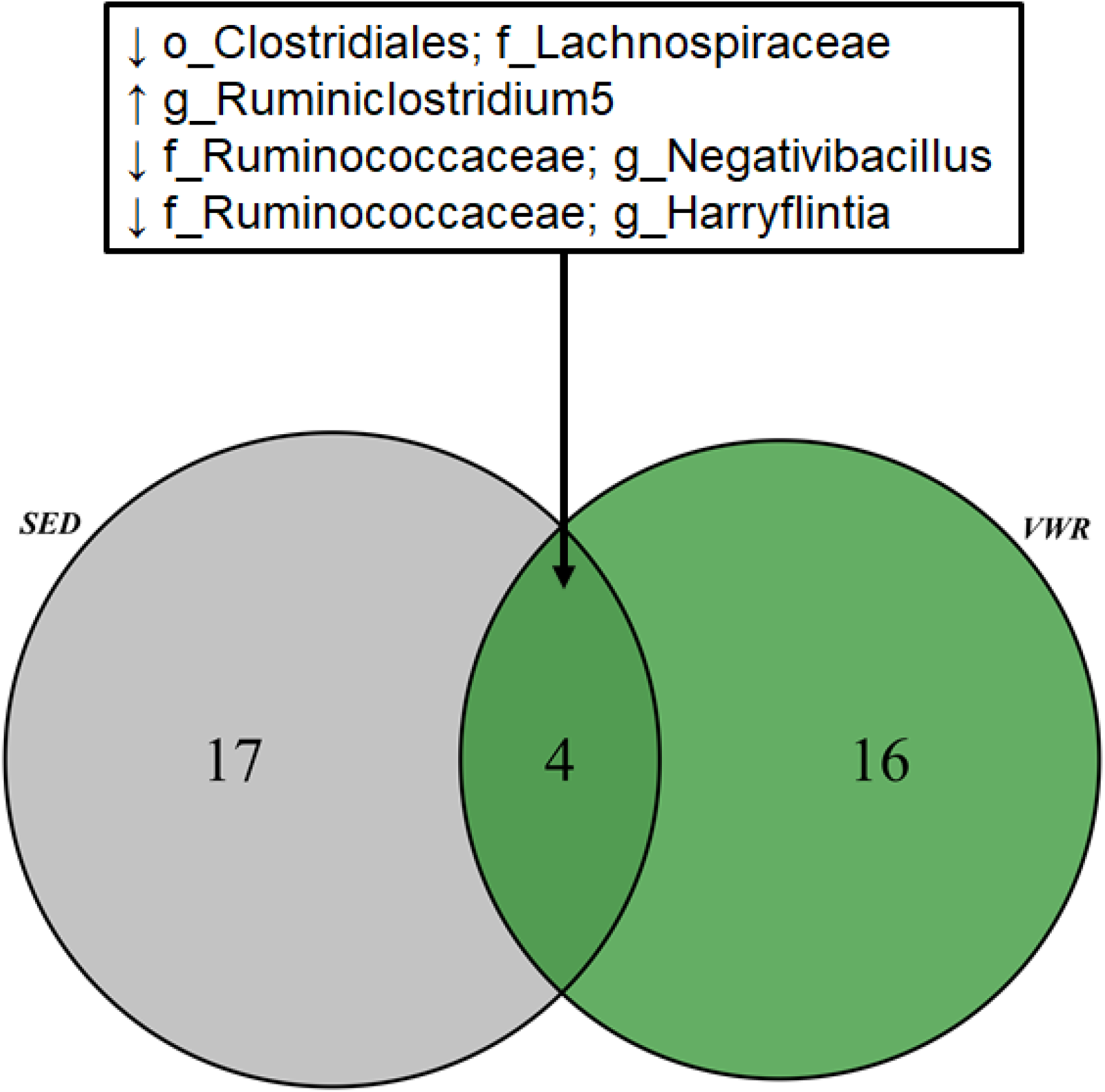
Venn diagram representing the number of unique ASVs detected to be differentially abundant across time in WT mice. The number in each circle represents the total number of ASVs that were statistically different at week 6. The overlapping region represents the number of significant ASVs that were changed in both groups. These 4 common taxa were changed in a similar fashion in both groups suggesting a possible age-driven effect, independent of wheel running. Arrows besides taxon name represent the direction of change at week 6 relative to week 0.

Predicted phenotypic traits: BugBase’s prediction of each community’s phenotypic traits suggest major differences between WT and *Muc2^-/-^* animals (Supplementary Material 5). Bacterial communities in *Muc2^-/-^* mice were composed of significantly higher abundances of Gram negative, aerobic, and facultative anaerobic bacteria with a higher potential for biofilm formation. Their communities also housed less bacteria with mobile elements and had an overall lowered tolerance for oxidative stress. At week 6, only VWR mice showed significant changes in their bacterial phenotypes compared to week 0. Their communities showed lower average abundances of mobile-containing (~12 %, P<0.05) and Gram-positive (~14 %, P<0.001) bacteria (Figure 8).

**Figure 8.**
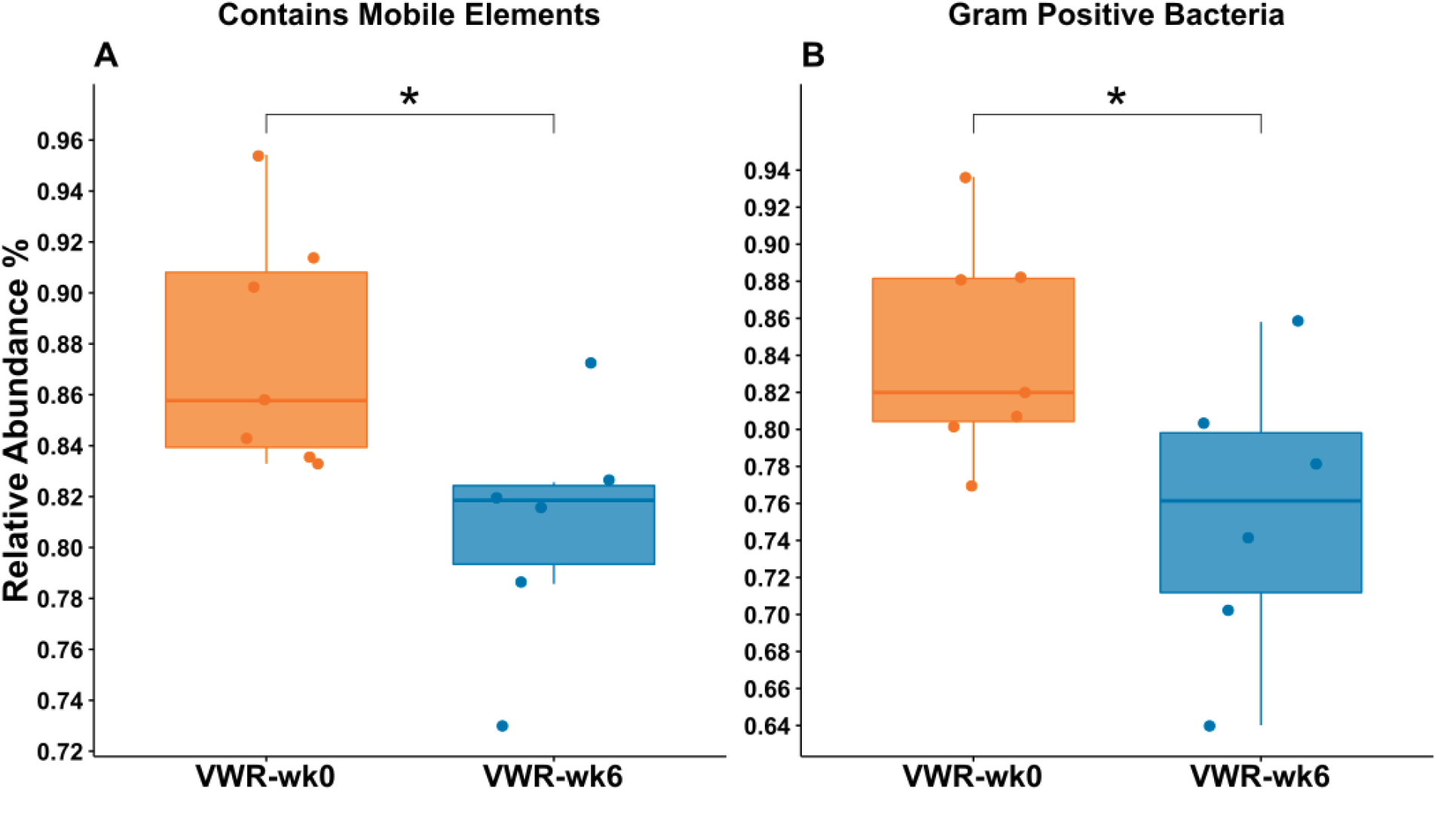
Change in predicted phenotypic traits of microbial communities in VWR mice following 6 weeks of wheel running. BugBase was used to predict the composition of microbial communities based on their bacterial traits. Only VWR mice showed a statistically significant change in overall microbial community traits of A) those containing a mobile element, and B) Gram positive bacteria, across time.

**Supplementary Material 5.**
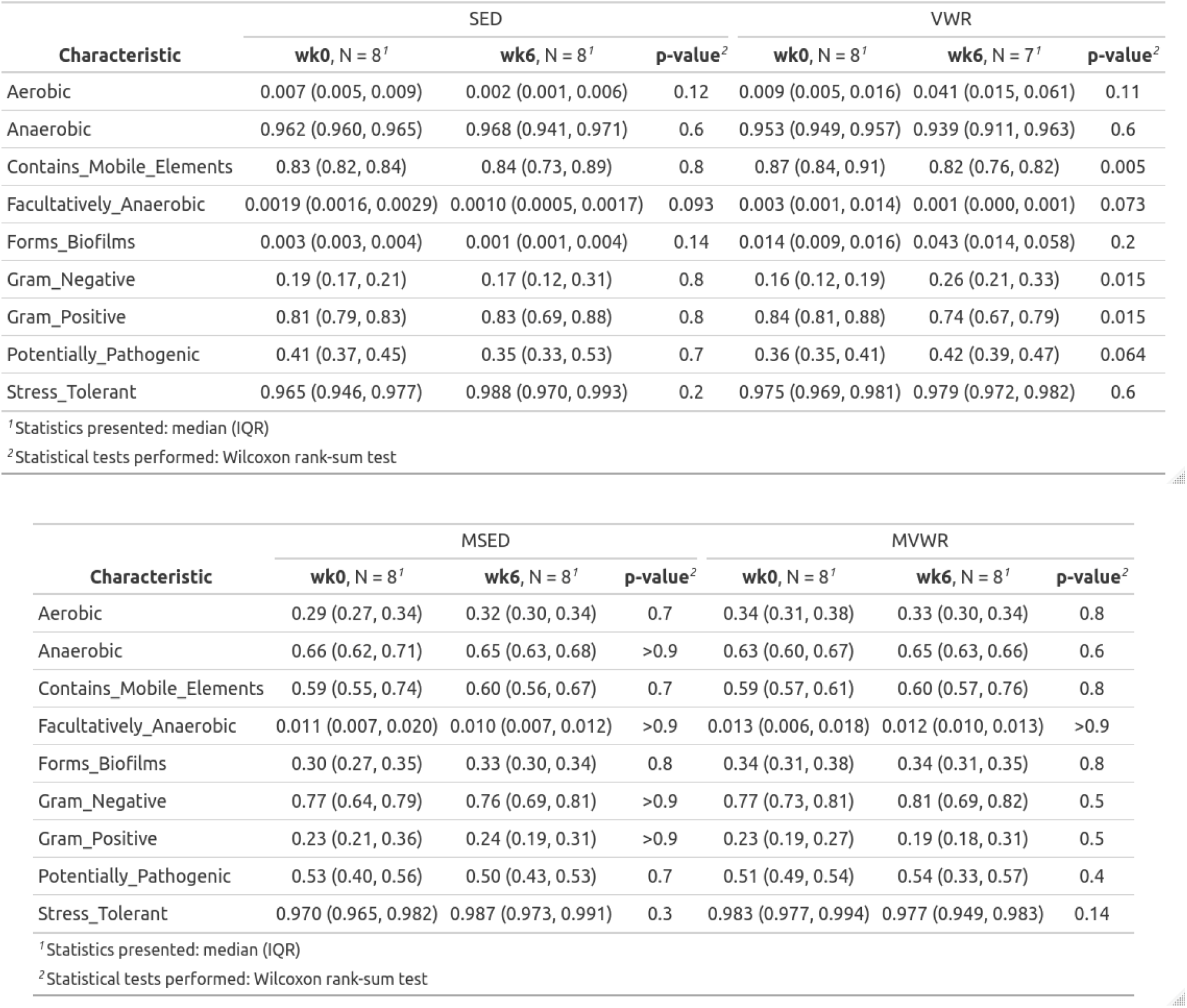
Summary statistics of predicted microbial traits. Bugbase was used to predict the phenotypic traits of microbial communities based on 16S marker genes. Values in the table are relative abundances of bacteria with the associated trait.

## Discussion

### *Muc2^-/-^* mice vastly differ from WT in their colonic cytokine, SCFA, and microbial profiles

*Muc2^-/-^* mice displayed clinical and histological symptoms of moderate colitis corresponding to the expected severity of this model at 11 weeks of age in our facilities. The colonic gene expression of inflammatory cytokine TNF-α, and the mucosal defense factor RELM-β, as well as antimicrobial peptide RegIII-γ were upregulated in *Muc2^-/-^* animals, as observed previously (15). Notably, the anti-inflammatory cytokine IL-10 was overexpressed in *Muc2^-/-^* compared to C57BL/6 WT. While in a healthy state, the expression of IL-10 may be associated with increased tolerance to inflammatory events, in *Muc2^-/-^* animals, this upregulation is essential in the host’s efforts at suppressing the excessive inflammation resulting from continuous exposure to bacterial ligands. Indeed, *Muc2^-/-^* + IL-10^-/-^ double knock-out mice show highly exacerbated colitis clinical signs (50) compared to deletion of either genes separately. The increase in IL-10 also has been previously observed in chemical models of colitis (9, 11). We further detected significant overexpression of CXCL9 in *Muc2^-/-^* animals. CXCL9 is a chemokine involved in regulating leukocyte trafficking, likely in response to exposure of bacterial ligands to host cells. CXCL9 overexpression also has been reported in IBD patients (51). Overall, the cytokine profile of *Muc2^-/-^* animals reflect those expected in human IBD.

*Muc2*^-/-^ mice born without a normal mucus layer house drastically less diverse and different bacterial communities than WT mice, as evident by the clear clustering of this group from WT animals in our PCA plots. Similar to patterns seen in IBD patients (52), or chemically-induced murine colitis (53, 54), *Muc2^-/-^* animals had an overall reduced α-diversity compared to their WT counterparts. The dominant taxa in WT mice were generally of the *Bacteroides* genus, Clostridiales order, and Lachnospiraceae family, while *Muc2^-/-^* animals were dominated by members of the Muribaculaceae family (formerly known as S24-7 (55)) and Akkermansia genus of the Verrucomicrobia phyla. While our taxonomic classifier was unable to confidently differentiate between the 2 sole species (Akkermansia muciniphila and Akkermansia glycaniphila) within this genus, it is reasonable to assume that the observed taxa were *A. muciniphila* as *A. glycanphilia* has, to date, only been isolated from python feces (56). *A. muciniphila* is perhaps the most surprising finding in this group as this species is known -and named-for its ability to degrade mucin, and is broadly considered as a beneficial bacterium in a variety of chronic diseases including IBD (57–59). The broader implications of this finding are beyond the scope of the present study. However, it does warrant the reassessment of the characterization of *A. muciniphila* as a mucin loving species to one that thrives in the absence of mucin. The bacterial phenotypic traits of *Muc2^-/-^* animals were predicted to be higher in abundances of Gram-positive, aerobic, and biofilm forming groups compared to WT mice. Lastly, the cecal SCFA of *Muc2^-/-^* mice were composed of significantly less butyrate and higher propionate concentrations compared to SED animals. The increased propionate levels in these animals is likely associated with the high abundances of *A. muciniphila*, a prominent propionate producer (60, 61). Overall, we found the *Muc2*^-/-^ model of colitis to capture many components of human IBD, especially those with impaired mucosal integrity.

### Wheel running in *Muc2^-/-^* mice does not reduce the severity of chronic colitis

Contrary to our primary hypothesis, we found that 6 weeks of wheel running in *Muc2^-/-^* mice did not improve the severity of clinical signs, histopathological scores, colonic expression of inflammatory cytokines, or abundances of cecal SCFAs, and did not alter the gut microbial composition in a consistent manner. These findings contrast others that show protective effects of VWR or forced treadmill running in chemically-induced models of colitis (8, 10, 11, 62). The fundamental difference between those studies and ours is in the model of colitis used. Previously, VWR was initiated in healthy animals prior to disease induction with chemical toxins, whereas in our study, wheel running is imposed over an existing disease state as a therapeutic intervention. This would suggest that PA prior to disease onset primes various components of intestinal health, enhancing its tolerance to injury. The effects of PA following disease-onset on the other hand, are either abolished or are overwhelmed by stronger disease signaling. It is also possible that the physiological benefits of PA depend on the presence of a healthy mucosal layer. This is supported by our findings that wheel running in WT but not *Muc2^-/-^* animals leads to significantly lower levels of pro-inflammatory colonic cytokines, increased anti-inflammatory IL-10, and increased levels of beneficial SCFAs. Given that UC patients typically have defective and thinning colonic mucosal layers, this would suggest that exercise prescription in these populations may have limited direct benefits on their intestinal health. However, the well-documented benefits of exercise are instituted across various other sites and systems of the body, which may indirectly result in improving primary and secondary disease symptoms through other pathways not accounted for in this experiment. For example, PA as a primary intervention has been associated with improved quality of life in IBD patients (63) and inversely correlated with loss of bone mass density, a common risk factor in this population (64, 65).

### VWR significantly attenuates pro-inflammatory, and upregulates anti-inflammatory cytokines in WT mice

Compared to SED animals, the wheel running mice showed lower levels of inflammatory cytokines TNF-α, IFN-γ, and TGF-β, all of which have been implicated in IBD (66). TNF-α is perhaps the most studied cytokine in relation to IBD as it plays a crucial role in innate and adaptive immunity and is directly involved in apoptotic processes in the intestines (67). It is found in significantly higher abundances in IBD patients (68) as well as murine colitis (69), making its regulation an obvious target for disease management. In fact, TNF-α inhibition using monoclonal antibodies is the most common target of biological therapies for moderate to severe IBD. The role of IFN-γ in colitis pathogenesis is less consistent across the literature, however its overproduction has been shown in CD (70, 71) and UC patients (72). In DSS-induced colitis models, neutralization antibodies against IFN-γ significantly reduced disease severity(73), while IFN-γ^-/-^ mice were completely protected from disease clinical signs (74). Anti-IFN-γ antibody treatments in human IBD are less effective however, with their efficacy dependent on baseline C-reactive protein levels (75), highlighting the need for treatment personalization. TGF-β is a pleiotropic cytokine that is ubiquitously produced by many cells and is involved in various immune functions including both anti- and pro-inflammatory actions. These include suppression of immune responses through recruitment of Tregs which in turn produce IL-10, but TGF-β can also elicit potent Th17 responses to combat extracellular bacteria (76). TGF-β is found in higher concentrations in intestines of IBD patients (77, 78), due to increased exposure of microbial ligands to host epithelial cells. Inversely, the attenuated levels of this cytokine in our VWR animals then may reflect a decrease in bacterial antigen exposure to the IEC suggesting reduced levels of host-microbe interactions in the mucosa. Alternatively, reduced TGF-β could also indicate reduced Treg activity in VWR mice, however, the increase in Treg derived IL-10 in these animals does not support this notion. IL-10 is an anti-inflammatory cytokine ubiquitously secreted by Tregs and is the primary driver of immunosuppressant actions in the intestines. Polymorphism in IL-10 promoters have been linked to IBD, making IL-10 supplementation a potential target for IBD therapy, however, clinical studies of IL-10 therapy to date have not been significantly effective (79). The significant increase in IL-10 in VWR mice suggests higher Treg activity which is associated with reduced inflammation. This is in agreement with others who showed a significant increase in murine intestinal IL-10 following treadmill running or swimming (80, 81). However, it is unclear whether this reflects a beneficial increase in anti-inflammatory events, or simply an adaptive response to changes in the microbial composition. Gram-negative bacteria preferentially stimulate IL-10 production and are associated with higher virulence due to increases in abundance of lipopolysaccharides bound to their cell walls (82). The higher expression of IL-10 in VWR animals then is likely correlated with increased abundance of Gram-negative bacteria observed in these mice. Further investigations are needed to determine the consequence of these changes. Taken together, the reduction of these pro-inflammatory cytokines and increase in anti-inflammatory IL-10 in VWR animals suggests a primed anti-inflammatory state in healthy WT but not diseased intestines, marking them as potentially important targets for prevention and reemission maintenance therapy.

### VWR significantly augments SCFAs content in WT but not *Muc2^-/-^* mice

SCFAs are metabolic by-products of bacterial fermentation of dietary fibers in the colon and are involved in various physiological processes of the host. Aberrant intestinal SCFAs content has been implicated in various diseases such as irritable bowel syndrome, cardiovascular disease, certain cancer types, and IBD (83–85). The most abundant of these, acetate, propionate, and butyrate, which make up over >95% of SCFA in humans (23), are markedly decreased in IBD patients (86), while their exogenous delivery can reduce inflammation *via* inhibition of TNF-α release from neutrophils (84, 87). Overall, increases in these SCFAs, especially butyrate, appear to positively influence IBD (88). We found an overall higher abundance of total cecal SCFAs, acetate, butyrate, and propionate in response to wheel running in WT but not *Muc2^-/-^* animals. This is in accordance with others showing higher butyrate concentrations in wheel running in rats (89), following exercise training in lean humans (30), and elite athletes (90). We’ve also previously observed a positive association between higher butyrate levels and VO2peak in healthy humans (16). The increase in these SCFAs may simply reflect higher energy demands of colonocytes which utilize SCFAs as their primary energy substrate. Interestingly, when we analyzed SCFAs content in relative abundances, we saw a significant increase in relative abundance of butyrate, but not acetate, or propionate. This suggests a preference in VWR animals for production of butyrate and its accompanying anti-inflammatory properties. These findings further support the patterns of anti-inflammatory priming we observe in these animals, contributing to an overall healthier intestinal environment following physical activity. The mechanisms behind PA-induced changes in SCFA are not known, however, given that SCFA are primarily produced by the intestinal microbiota, it is highly likely that changes in SCFA are linked to the observed changes in the microbiome. SCFA affects microbiota dynamics as they are directly involved in chemical balance and pH regulation of the intestines (91) and in turn the microbiota can also affect SCFA production and use, establishing a bidirectional affiliation.

### Wheel running has limited but significant effects on the intestinal bacterial composition of WT but not *Muc2^-/-^* mice

In this study, neither time nor wheel running had any effect on any alpha diversity metrics measured. The effects of PA on alpha diversity is not consistent within the literature. For example, the findings here are in contrast to our own previous observations in healthy humans that showed a significant correlation between alpha diversity and cardiorespiratory fitness (16). Others have also reported that elite athletes have higher alpha diversity than sedentary controls (92), or that exercise training in mice leads to increased Shannon diversity (93, 94). However, in agreement with the current experiment, PA has been shown to have no effect on alpha diversity in mice (29, 95, 96), rats (97), and humans (30, 98). The reasons for these discrepancies are not clear, though multiple factors such as differences in animal vendors and facilities, DNA extraction methods and sequencing, bioinformatics analysis, and statistical testing methods, are likely involved. Additionally, one particularly important consideration in comparing animal models of PA is the total volume of activity performed. For example, across the aforementioned studies utilizing wheel running in mice, we noticed a wide range (~ 2.5-10 km) of average daily running distances reported. Different volumes of PA are likely to elicit different physiological responses which can extend to the microbiome.

Comparisons of the across-samples diversity (beta diversity) in *Muc2^-/-^* animals showed no patterns of change as a function of time or wheel running. In WT animals, however, the change in Aitchison distances between week 6 and 0 were significantly different between VWR and SED groups, indicating both time and wheel running as important factors in the observed shift in community composition. The magnitude of this change across time between the groups however was not different.

Univariate analyses of individual taxa in *Muc2^-/-^* animals also showed no significant changes in any microbial clades across either groups. A lack of significant change in these animals suggests that the presence of a healthy mucosal layer is required to mediate PA-induced changes in community composition in the colon. In WT animals, the relative abundances of over 20 taxa in each group were significantly different by week 6 (Figure 7), with only four of those taxa common to both VWR and SED groups. The changing taxa in either group belonged primarily to the Ruminococcaceae and Lachnospiraceae families, with some species increasing while others decreased. Notably, In SED animals, 5 species from the Bacteroides genus were decreased by week 6, while no changes in this genus were detected in VWR animals. Species within the Bacteroides genus are Gram-negative, obligate anaerobes that are among the most abundant found within the mammalian intestine and carry important functions such breaking down of complex glycans, refining the gut environment by reducing intracellular oxygen levels, and preventing the colonization of opportunistic pathogens (99). Bacteroides spp. are also one of the primary propionate producers in the mammalian gut (100), therefore, the observed reduction in members of this group in SED animals may in part explain their lower propionate levels compared to VWR mice. Given that we were unable to classify these ASVs to the species level, it is difficult to speculate further on the biological implications of these observations. Perhaps the most studied species in this group, *Bacteroides fragilis*, has been shown to be protective against DSS colitis in mice by stimulating IL-10 expression (101, 102). Given that we also observed significantly lower IL-10 expression in SED mice, it is reasonable to speculate that at least one of the unclassified Bacteroides ASVs in this group is *B. fragilis* and linked to the relatively lower expression of this protective cytokine in those animals.

Analyzing the bacterial consortia based on their predicted phenotypic traits revealed additional information regarding the effect of wheel running on the overall community. Following wheel running, WT but not *Muc2^-/-^* mice, had significant reduction (~ 14%) in total abundance of Gram-positive bacteria. This is supported by the observed decreases in several members of the Gram-positive Ruminococcaceae in these mice, as well as the attenuated expression of RegIII-γ, an antimicrobial peptide that specifically targets the surface peptidoglycan layer of Gram-positive bacteria. The implications of this phenotypic shift in microbiota of healthy individuals is not known, but may provide a clue for understanding the adaptations of the intestinal environment to the physiological stresses of PA. Furthermore, mirroring the shift in Gram-positive phenotype was the decreased relative abundances of bacteria containing mobile elements. These refer to microevolutionary processes such as transposons i.e. segments of DNA with the ability to move locations within the genome, and bacterial plasmids which are involved in horizontal gene transfer. These events are typically associated with sharing of virulence factors between bacterial cells and increased resistance to antibiotics. The higher abundances of mobile elements in bacteria from these mice is likely not indicative of antibiotic-resistance but rather associated with higher abundances of Gram-negative bacteria representing more mobile-elements. The results of these predictions should be interpreted with caution however, as these mobile elements can rapidly become population specific within an individual thus precluding inference across similar experimental groups (103).

## Summary

In contrast to our hypothesis we found that 6 weeks of wheel running did not ameliorate any clinical signs of colitis in *Muc2*^-/-^ animals, nor did it influence any components of the intestinal environment such as expression of various cytokines and production of SCFA. Wheel running in healthy WT C57BL/6 mice on the other hand, imposed various physiological effects on the gut, including downregulation of pro-inflammatory and upregulation of anti-inflammatory cytokine gene expression, and increased concentration of total SCFAs including butyrate, acetate, and propionate. Wheel running further led to a shift in bacterial community structure corresponding to higher abundances of Gram-negative bacteria. As these physiological changes have been associated with protection against chronic inflammatory diseases in humans such as IBD, we conclude that PA prior to disease onset can prime the intestines, enhancing their tolerance to inflammation. These benefits however are lost when PA is imposed in the absence of a healthy mucosal layer. Overall, the findings here suggest that PA in healthy individuals may be an important preventative medicine against intestinal diseases such as IBD.

**Supplementary Material 6.**
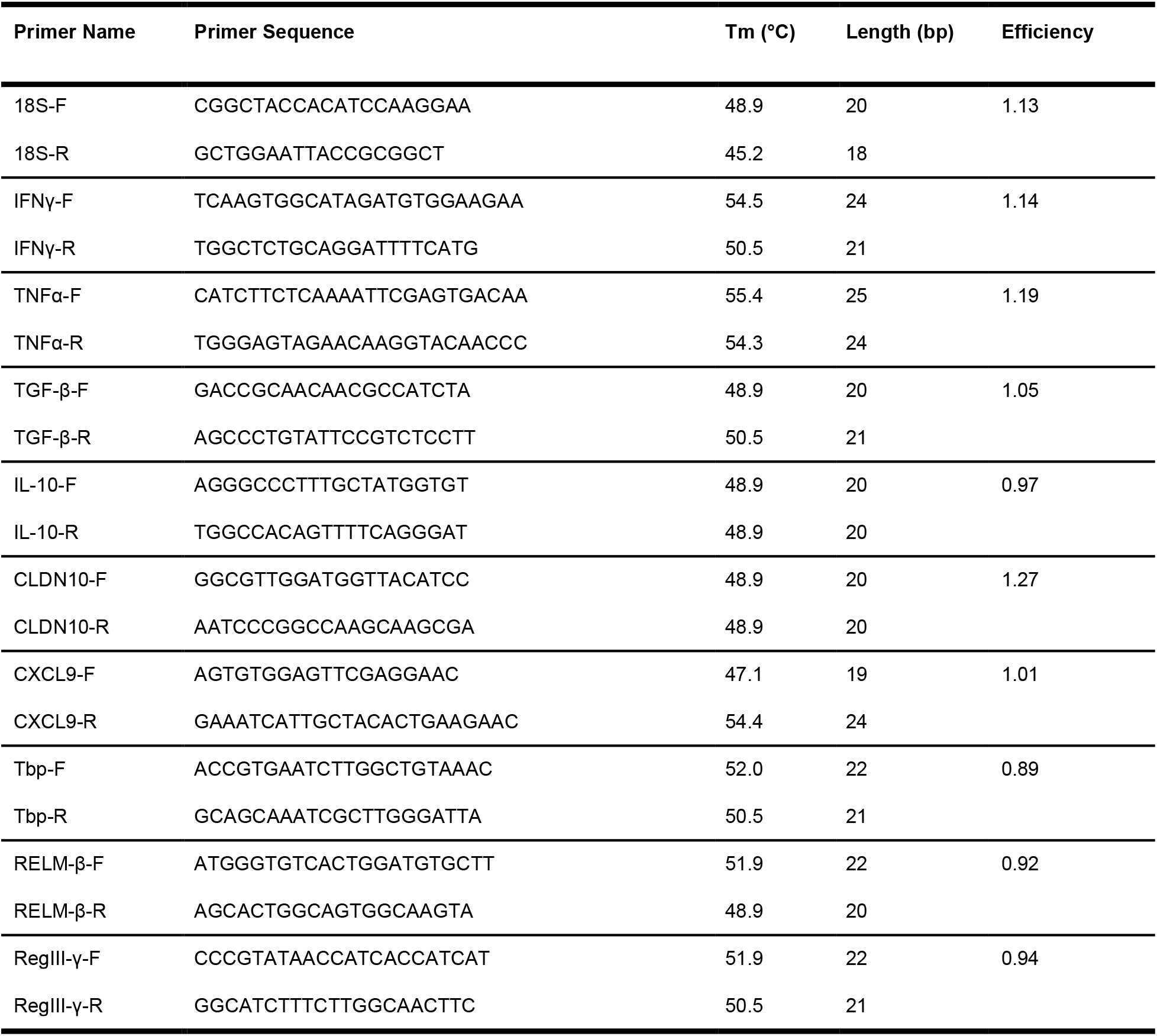
List of RT-qPCR primers used. Tm refers to the primer melting temperature. Length (bp) refers to primer length in base pairs. Primer names ending with ‘-F’ or ‘-R’ refer to forward and reverse primers, respectively.

## Availability of data

All data and codes used for all aspects of this manuscript, including all metadata, histology figures, and raw sequence files are currently available publicly at https://osf.io/9awgd/

All used software packages, versions, and parameters from the QIIME 2 software are available under the ‘provenance’ tab of the accompanying qiime2 zip artifact (.qza) files. These files can be viewed locally or on a web browser at https://view.qiime2.org/.

## Author contributions

**Conceptualization:** ME, DLG; **Data curation:** ME; **Formal Analysis:** ME, JP; **Funding acquisition**; DLG; **Investigation:** ME, DWM, CQ, AG, SKG, JAB; **Methodology:** ME, DLG; **Project administration:** ME, DLG; **Resources:** DLG; **Software:** ME, JP; **Supervision:** DLG; **Validation:** ME, JP; **Visualization:** ME; **Writing – original draft:** ME; **Writing – review & editing:** ME, DWM, CQ, JP, JB, SG, DLG

